# CD28 Shapes T Cell Receptor Signaling by Regulating ZAP70 Activation and Lck Dynamics

**DOI:** 10.1101/2024.06.27.601067

**Authors:** Kumarkrishna Raychaudhuri, Rohita Rangu, Alison Ma, Neriah Alvinez, Andy D. Tran, Sandeep Pallikkuth, Katherine M. McIntire, Joseph A. Garvey, Jason Yi, Lawrence E. Samelson

**Author notes:** Corresponding author: Laboratory of Cellular and Molecular Biology, Center for Cancer Research, National Institutes of Health, Bethesda, MD, 20892, USA.

## Abstract

T cell activation requires T cell receptor (TCR) engagement, which initiates a series of proximal events including tyrosine phosphorylation of the CD3 and TCRζ chains, recruitment, and activation of the protein tyrosine kinases Lck and ZAP70, followed by recruitment of adapter and signaling proteins. CD28 co-stimulation is also required to generate a functional immune response. Currently we lack a full understanding of the molecular mechanism of CD28 activation. TCR microclusters (MC) are submicron-sized molecular condensates and basic signaling units that form immediately after TCR ligation. Our results show that CD28 co-stimulation specifically accelerated recruitment of ZAP70 to the TCRζ chain in MCs and increased ZAP70 activation. This CD28-mediated acceleration of ZAP70 recruitment was driven by enhanced Lck recruitment to the MCs. A greater spatial separation between active and inactive species of Lck was also observed in the MCs as a consequence of CD28 co-stimulation. These results suggest that CD28 co-stimulation may lower the TCR activation threshold by enhancing the activated form of Lck in the TCR MCs.

## Introduction

T lymphocytes are central to the adaptive immune response. To become activated, the T cell antigen receptor (TCR) engages with antigenic peptide loaded on major histocompatibility complex-encoded proteins (p-MHC) on an antigen presenting cell (APC). The TCR-p-MHC interaction leads to phosphorylation of the immunotyrosine based activation motifs (ITAMs) of the CD3 δ, ε, γ and TCRζ chains by the Src-family protein tyrosine kinase Lck. Fully phosphorylated ITAM motifs recruit the Syk-family kinase, ZAP70, which is then phosphorylated and activated by Lck. Activated ZAP70 phosphorylates and activates several downstream signaling and adapter proteins such as LAT, GRB2, SLP76, ADAP, NCK, VAV1, PLCγ and c-Cbl^1–3^. LAT is a scaffold on which most of the signaling and adaptor proteins assemble and subsequently become activated. TCR engagement leads to these molecular events in a sequential and non-stochastic manner resulting rapidly in formation of MCs. MCs are phase-separated molecular condensates found at the interface of the T cell and the activation surface. As aggregates of multiple signaling, adapter, and receptor proteins involved in TCR activation, MCs function as engines of T cell activation, and they are critical to formation of the immune synapse^4–6^. High resolution spatio-temporal studies of molecular events within these MCs revealed important proximal events of TCR activation^3^. It was observed that TCR and ZAP70 localize to form a receptor domain, which is spatially separated from a signaling domain comprised of LAT, GRB2, SLP76, NCK, ADAP, VAV and PLCγ1. We also observed temperature and calcium-dependent distinct kinetic lags between recruitment of receptor domain proteins, and between receptor and signaling domain proteins.

TCR-pMHC ligation alone results in an unresponsive state in the T cell known as anergy^7,8^. Interaction of CD28 on the T cells with its ligands, B7.1 (CD80) or B7.2 (CD86), on the APC provides the crucial second signal required to generate a functional immune response in a naïve T cell. CD28 is a constitutively expressed co-stimulatory molecule with an extra-cellular domain characteristic of the immunoglobulin family^9,10^. CD28 and TCR signaling have been shown to have a synergistic and cooperative relationship. CD28 enhances TCR efficiency, while sustained TCR signaling is shown to rapidly induce reorientation in the cytoplasmic domain of CD28 to increase the latter’s avidity for its ligand, CD80^11,12^. Because of such intricate functional and structural cooperativity between TCR and CD28 co-stimulation, one could imagine a proximal role for CD28 in regulating TCR signaling events. Moreover, the membrane proximal tyrosine and the membrane-distal proline-rich domains in the cytoplasmic tail of CD28 have been shown to interact with multiple proximal kinases and signaling proteins such as Lck, Itk, PKCθ, FLNA and GRB2^10,13,14^, suggesting a proximal role for CD28 in TCR signaling.

The TCR is unique in its ability to generate an immune response by responding to a surprisingly low number of weak affinity antigens on APCs. The consequence of the ability of the TCR to discriminate low affinity self vs non-self p-MHC interactions is addressed in the “Kinetic Proof-Reading Model”, which proposes that differences in receptor affinity for the pMHC ligand directly translate into differences in receptor dwell time and occupancy^15,16^. Successful interactions determine subsequent transduction of signal through contingent downstream steps leading to the required formation of protein-protein interactions and protein complexes. CD28 likely plays a crucial role in enhancing TCR-mediated signaling; however, our understanding of how such activity is carried out by CD28 at the proximal level is far from complete.

In this study we showed that CD28 co-stimulation plays an important role in determining the kinetics of a proximal step in TCR signaling. We observed that CD28 co-stimulation accelerates recruitment of ZAP70 to the TCRζ chain in the MCs, as revealed by a decreased kinetic lag between engagement of TCRζ and the recruitment of ZAP70. We also found that CD28 co-stimulation markedly changes Lck dynamics in T cells resulting in Lck enrichment within MCs and a concomitant increase in co-localization of Lck with ZAP70 and TCRζ. We also observed that CD28 co-stimulation results in greater spatial separation between activated and inhibited species of Lck, thereby increasing the possibility that there are localized Lck activation sites in MCs.

Based on our results, we hypothesize that CD28 co-stimulation lowers the TCR activation threshold by recruiting the activated form of Lck into the TCR MCs. Our results reveal a previously unknown mechanism of function for CD28 co-stimulation. These findings have the potential to be exploited in various aspects of immunotherapies such as development of more efficient chimeric antigen receptor constructs.

## Results

### CD28 co-stimulation accelerates recruitment of ZAP70 to TCRζ in the TCR MCs

In this study we describe the kinetic relationships between the recruitments of the TCRζ chain, the ZAP70 kinase and distal signaling proteins into microclusters (MC) in response to anti-CD3ε mediated TCR engagement^3^. We compare the kinetics of that stimulation with that induced by anti-CD3ε engagement with simultaneous CD28 co-stimulation. For these comparative studies we employed TIRF microscopy to elucidate the kinetic relationships in recruitment of these proteins in high spatio-temporal resolution.

To induce CD28 co-stimulation, we used anti-CD28 antibodies or natural ligands, B7.1 and B7.2. These natural ligands are recombinant proteins containing the extracellular region of B7.1 or B7.2 fused to the Fc portion of human IgG1. For TCR stimulation we used anti-CD3ε antibody^3^. Antibodies and the natural ligands were used alone or in combination to create different stimulatory conditions for T cells. Anti-CD3 stimulation alone resulted in induction of ZAP70 MCs but failed to recruit CD28 when Jurkat T cells expressing ZAP70-Emerald, and CD28-Apple were activated. Anti-CD28, B7.1 or B7.2 stimulation alone specifically recruited CD28 into MCs, but failed to recruit ZAP70 into those clusters. These results confirm the specificity and mutual exclusiveness we can achieve in our simulation of TCR triggering and CD28 co-stimulation. Importantly, when we combined anti-TCR and anti-CD28 stimulation by addition of anti-CD28 or either B7.1 or B7.2 we could trigger both TCR stimulation as well as CD28 co-stimulation as evident from formation of MCs containing both ZAP70 and CD28 (Supp. Fig. 1).

To determine the changes in the kinetic relationship between TCRζ and ZAP70 when stimulated with either TCR stimulation alone or in combination with CD28 co-stimulation, Jurkat T cells and mouse primary T cells expressing TCRζ-Emerald, and ZAP70-Apple were activated under those two conditions. As expected from our previous studies, TIRF time-lapse images confirmed a delay in recruitment of ZAP70 to TCRζ when cells were subjected to TCR stimulation alone (Fig 1a-c)^3^. We demonstrated this effect by tracking one microcluster at a time and showing the kinetics of recruitment of TCRζ and ZAP70 separately (Fig 1b). Normalized fluorescent intensity of the accumulation of these two proteins was graphed (Fig. 1c). Surprisingly when TCR stimulation was combined with CD28 co-stimulation we observed a faster recruitment of ZAP70 to TCRζ in microclusters (Fig. 1d-f). To compare the effects on large numbers of microclusters we determined the half-maximal level of normalized fluorescent intensity and plotted the kinetic lag between TCRζ and ZAP70 for each cluster (Fig. 1g). Measurement of the kinetic lag between TCRζ and ZAP70 in a large number of MCs in Jurkat T cells confirmed a significant decrease in kinetic lag from an average of ∼25s with anti-CD3ε alone to ∼20s with combined anti-CD3ε and anti-CD28 co-stimulation.

**Figure 1:**
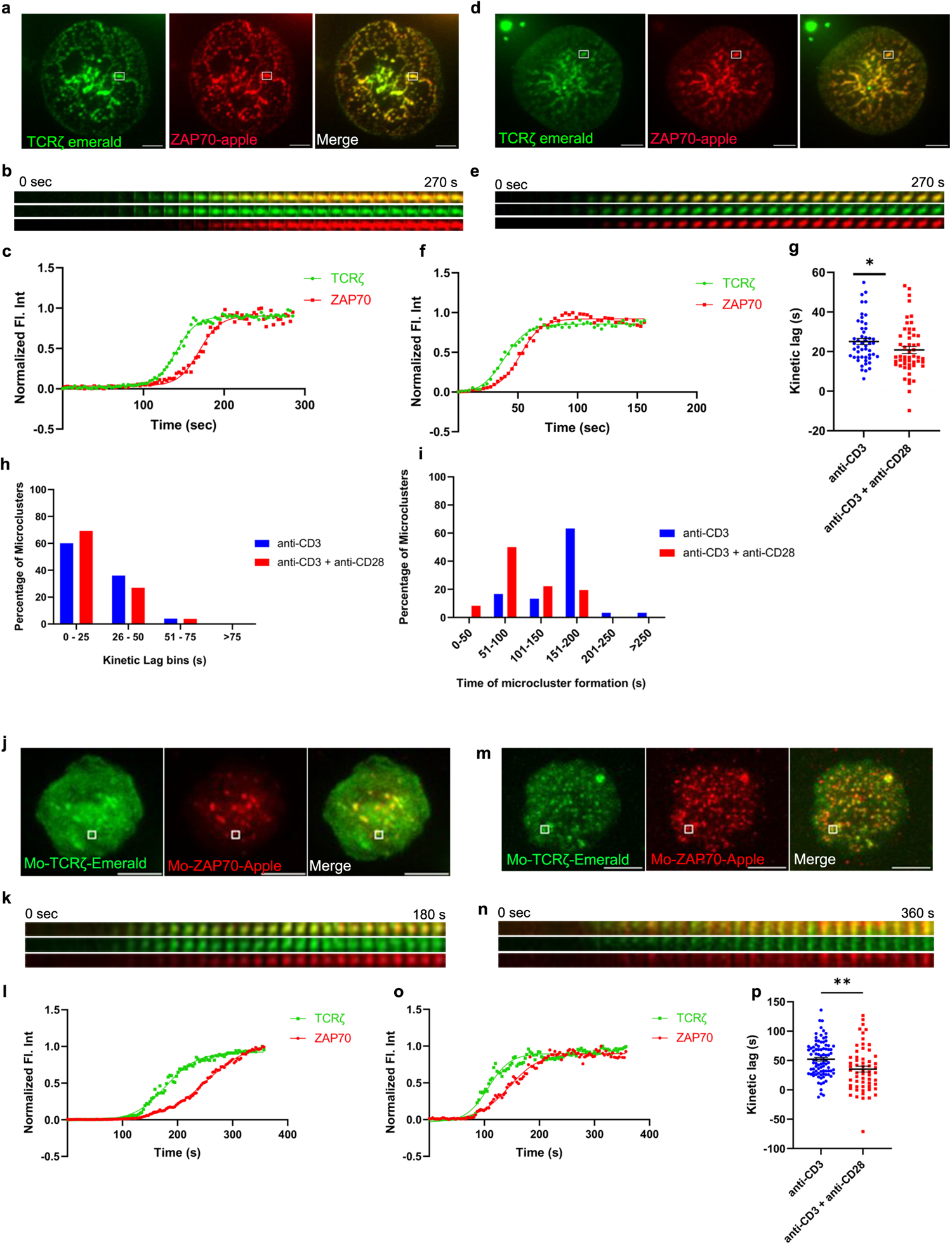
CD28 co-stimulation accelerates recruitment of ZAP70 to TCRζ in the microclusters of induced T cells. Jurkat T cells (**a**-**i**) or mouse primary T cells (**j**-**p**) were transfected to express TCRζ-Emerald (green) and ZAP70-Apple (red) and were activated on coverslips coated with either anti-CD3 alone (**a** and **j**) (Jurkat: *n =* 50 microclusters (MC), more than 7 (>7) cells; mouse primary T cells: *n =* 89 MC, >8 cells) or with anti-CD3 + anti-CD28 (**d** and **m**) (Jurkat: *n =* 52 MC, >7 cells; Mo-primary T cells: *n =* 59 MC, >5 cells) antibodies. 120 time-lapse images were acquired every 3s at 21°C using TIRF microscopy. **a, d** and **j, m**— Sum of all the image stacks for the individual time points (Max intensity projection) of representative Jurkat T cells or mouse primary T cells stimulated as indicated above. **b, e** and **k, n**— Time-lapse montage of representative microclusters from a, d and j, m, as indicated by white boxes. **c, f** and **l, o**— Representative relative intensity plot of the individual microclusters, as defined in the materials and methods section, indicated in **a, d** and **j, m,** respectively. **g** and **p**— Kinetic lags, as defined in the materials and methods section, measured between TCRζ and ZAP70 with indicated stimulatory antibodies. Each point represents the time lag, as defined in the materials and methods section, at half-max intensity of an individual microcluster. **h**— Distribution of average kinetic lags across specified time bins. **i**— Distribution of percentage of microclusters formed at specified time intervals of the first kinetic lag bin, as defined in the materials and methods section, (0-25s) in **h**. Data presented as mean ± SEM. Populations were analyzed using Mann-Whitney (Jurkat data) and Welch’s (mouse primary T cells) t-tests. \**p* < 0.05, \*\**p* < 0.01. Scale bar: 5μm.

A similar acceleration in recruitment of ZAP70 was also observed in a primary mouse T cell microcluster in response to anti-CD3ε and anti-CD28 co-stimulation as compared to stimulation with anti-CD3ε stimulation alone (Fig. 1j-l and 1m-o). Analysis of the accelerated recruitment of ZAP-70 induced by co-stimulation in multiple microclusters, as described above, was more impressive with an average TCRζ to ZAP70 gap decreasing from ∼52s to ∼35s in microclusters from these mouse primary T cells (Fig. 1g and p).

To exclude any anti-CD28 antibody-specific artifact we used combinations of anti-CD3ε and anti-CD43 antibodies to activate Jurkat T cells. Multiple microclusters were evaluated in cells activated by anti-CD3ε alone, by anti-CD3ε and anti-CD28 or by anti-CD3ε and anti-CD43 Measurement of the kinetic lag in this control group showed no change and a mean kinetic lag of ∼26s was observed between recruitment of TCRζ and ZAP70 which is comparable to anti-CD3ε stimulation alone (Supp. Fig. 2a).

The populational kinetics lag data exhibited a large distribution of time-lags and prompted us to group MCs from an experiment into several kinetic lag bins. In the CD28 co-stimulation group we observed a dramatic increase in the percentage of MCs falling into the early kinetic lag bin defined as 0s to 25s (Fig. 1h). Further analysis of the 0-25s kinetic lag bin revealed that MCs from cells which received co-stimulation tend to form early compared to MCs of their counterparts from cells that received TCR stimulation alone (Fig. 1i). These results indicate that CD28-co-stimulation results in more clusters in which the recruitment of ZAP70 is faster.

### CD28 co-stimulation has a less pronounced effect in regulating the kinetic lag between ZAP70 and signaling domain proteins

Receptor and ZAP70 clustering downstream of TCR ligation is rapidly followed by MC recruitment of signaling domain proteins, such as LAT, GRB2, SLP76, GADS, and ADAP. To investigate if CD28 co-stimulation affects the kinetics of recruitment of these proteins to the TCRζ and ZAP70 receptor domain proteins, we investigated the kinetics of recruitment of GRB2, SLP76, and GADS to the MC in relation to ZAP70. The time gap between recruitment of ZAP70 and GRB2 remained unchanged in Jurkat cells when they were stimulated with anti-TCR without or with anti-CD28 (Fig 2a-c and 2d-f). An individual microcluster was tracked in Fig. 2b and 2c for anti-TCR stimulation and in Fib. 2e and 2f for anti-TCR and anti-CD28 stimulation. The mean kinetic lag between recruitment of ZAP70 and recruitment of GRB2 in Jurkat cells was measured at ∼25s, which is consistent with our previous results from our laboratory^3^. CD28 co-stimulation did not change the kinetic lag between ZAP70 and GRB2 (∼24s) in Jurkat cells (Fig. 2g). However, in murine T cells, GRB2 recruitment to ZAP70 was significantly accelerated when cells were TCR stimulated in presence of CD28 co-stimulation (Fig. 2h-j and 2k-m). We observed a decrease in the mean kinetic lag from ∼23s to ∼9s between ZAP70 and GRB2 when stimulated with anti-TCR and anti-CD28 stimulation compared to anti-TCR stimulation alone (Fig 2n). Possible explanations for this striking difference in the ZAP-70-Grb2 gap between Jurkat and murine T cells are proposed in the Discussion. For the remainder of this work, we restricted our focus on the contribution of CD28 co-stimulation to the most upstream event following TCR engagement.

**Figure 2:**
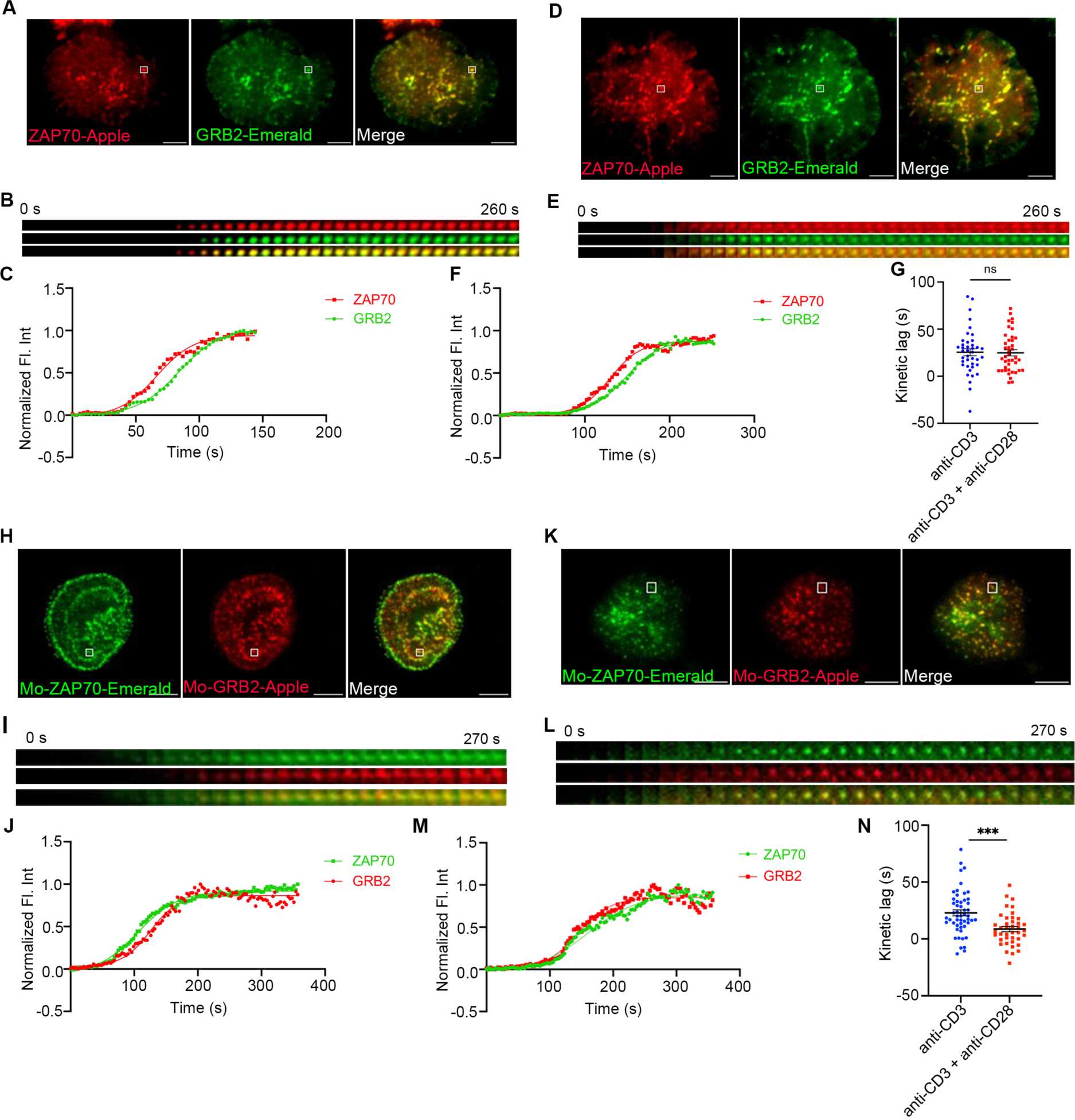
The effect of CD28 co-stimulation is less pronounced in regulating the kinetic lag between ZAP70 and signaling domain proteins. Jurkat T cells (**a**-**g**) or mouse primary T cells (**hn**) were transfected to express ZAP70-Apple (red) and GRB2-Emerald (green) or ZAP70-Emerald (green) and GRB2-Apple (red), respectively. Cells were activated on coverslips coated with either anti-CD3 alone (**a** and **h**) (Jurkat: *n =* 40 MC, >8 cells; Mo-primary T cells: *n =* 52 MC, >8 cells) or with anti-CD3 + anti-CD28 (**d** and **k**) (Jurkat: *n =* 41 MC, >11 cells; Mo-primary T cells: *n =* 40 MC, >6 cells) antibodies. 120 time-lapse images were acquired every 3s at 21°C using TIRF microscope. **a, d** and **h, k**— Max intensity projection of representative Jurkat T cells or mouse primary T cells stimulated as indicated above. **b, e** and **i, l**— Time-lapse montage of representative microclusters from a, d and h, k respectively, as indicated. **c, f** and **j, m**— Representative relative intensity plot of the individual microclusters indicated by white boxes in a, d and h, k respectively. **g** and **n**— Kinetic lags measured between ZAP70 and GRB2 with indicated stimulatory antibodies. Data presented as mean ± SEM. Populations were analyzed using Student’s t-tests. \**p* < 0.05, \*\**p* < 0.01, \*\*\**p*<0.001. Scale bar: 5μm.

Stimulation of Jurkat cells with the combination of anti-CD3ε and anti-CD43 yielded a kinetic lag between ZAP70-GRB2 and ZAP70-SLP76 similar to anti-CD3ε stimulation alone, ruling out any nonspecific effect of ant-CD28 antibody (Supp. Fig. 3a and 3d). Unlike our observations of the TCRζ to ZAP70 gap we did not observe that MCs from the anti-TCR plus anti-CD28 group preferentially segregated into a short kinetic lag cohort when the populational data was grouped into several kinetic lag bins (Supp. Fig. 3b). MCs from cells co-stimulated by anti-CD28 from the short kinetic lag cohort did not exhibit any change in the time of their formation when compared to MCs from the anti-TCR stimulation group alone (Supp. Fig. 3c). We also investigated the kinetic lag from ZAP70 to SLP76 and ZAP70 to GADS in Jurkat cells but did not observe any acceleration in recruitment of SLP76 or GADS to ZAP70 (Supp. Fig. 3d and 3f). Also, CD28 co-stimulation did not skew the MC distribution to the shorter kinetic lag when compared to anti-TCR stimulation alone in cells where ZAP70-SLP76 and ZAP70-GADS kinetics were measured (Supp. Fig. 3e and 3g). These results suggest that CD28 co-stimulation predominantly modulates TCR signaling events as early as recruitment of the kinase ZAP70 in Jurkat cells. Recruitment of signaling domain proteins, such as GRB2, SLP76 and GADS are further downstream effects of CD28 costimulation and may require secondary signaling events.

### CD28 co-stimulation accelerates recruitment of E3 ligase, cCbl to TCRζ but not to ZAP70

In a previous study we showed that c-Cbl is recruited to the MC after Grb2 and other signaling proteins^3^. c-Cbl is known to ubiquitinate ZAP-70 and LAT which results in their internalization and degradation, triggering dissociation of the MC signaling complex^17–19^. Therefore, we used c-Cbl recruitment as an early readout of dissociation of signaling complexes in the MC. We simultaneously mapped the kinetic relationship between TCRζ, ZAP70, and c-Cbl, a “three-protein expression system,” as they were concurrently expressed and sequentially recruited to the MC. In these experiments we again observed a significant acceleration in recruitment of ZAP70 to TCRζ in response to the combination of TCR and CD28 co-stimulation compared to TCR stimulation alone (Fig. 3a-g). There is a delay in recruitment of c-Cbl to ZAP70 in Jurkat cells when stimulated with anti-TCR antibody, which is consistent with our previously published results (Fig. 3a-c)^3^. Combining TCR stimulation and CD28 co-stimulation did not accelerate recruitment of c-Cbl to ZAP70, and no significant change in the mean kinetic lag between ZAP70 and c-Cbl with or without CD28 co-stimulation was observed (Fig. 3a-f, and h). As would be expected from combining these two lags, the kinetic lag between TCRζ and c-Cbl showed a significant decrease when cells received CD28 co-stimulation along with TCR stimulation (Fig. 1a-f, and i). These results suggest that c-Cbl recruitment to the MC is accelerated in the presence of CD28 co-stimulation and dissociation of the signaling complex may occur faster in comparison to TCR stimulation alone. A shorter life span of the signaling complex suggests that CD28 co-stimulation improves the efficiency of the molecular interactions in the signaling complex and accelerates translation of those interactions to cellular signals prior to complex dissociation. Accelerated recruitment of ZAP70 is sufficient to relay the effects downstream of CD28 co-stimulation.

**Figure 3:**
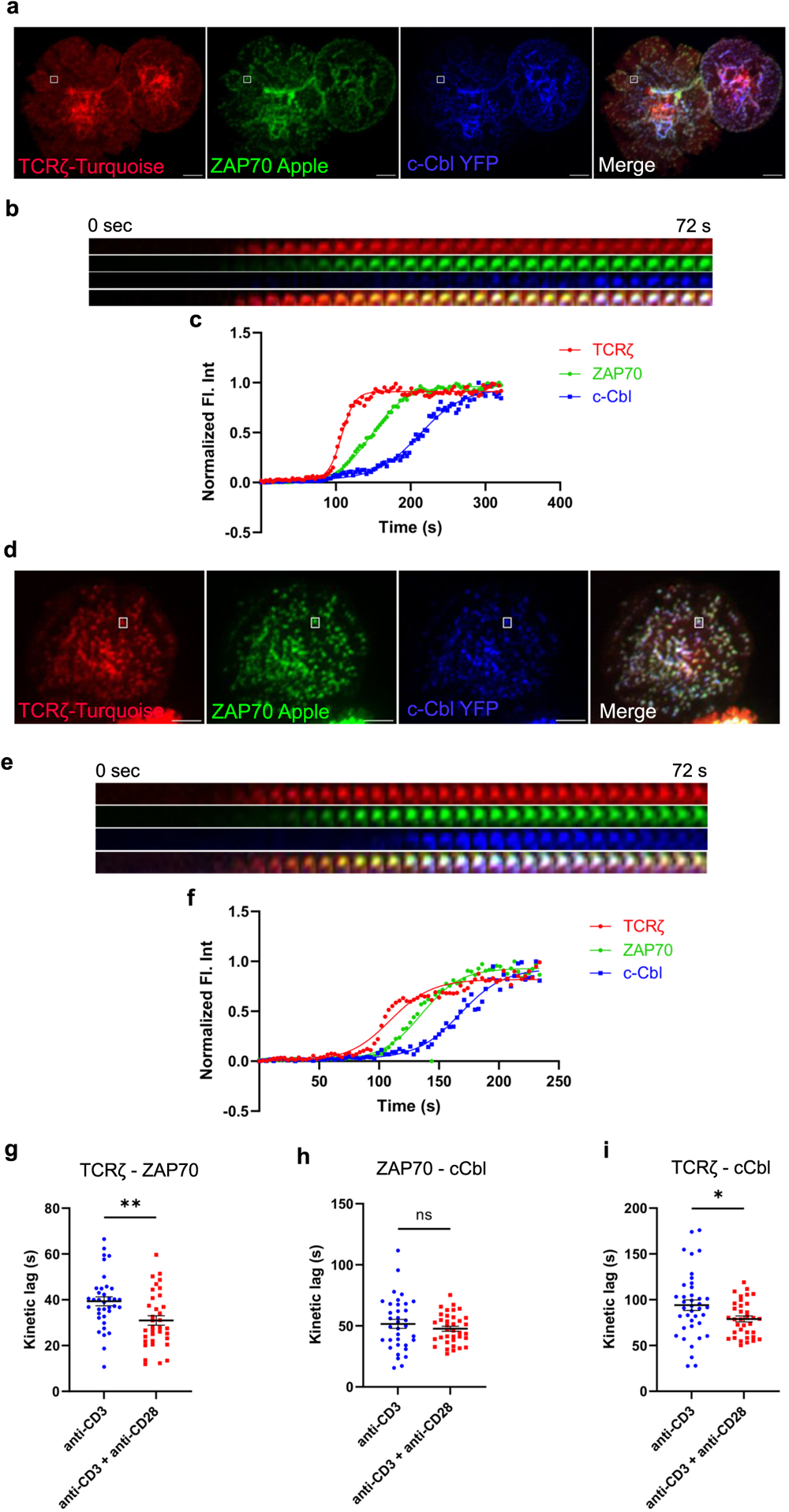
Recruitment of E3 ligase c-Cbl to the TCRζ but not to ZAP70 is accelerated by CD28 co-stimulation. Jurkat T cells were transfected to express TCRζ-Turquoise (red), ZAP70-Apple (green) and c-Cbl-YFP (blue). Cells were activated on coverslips coated with either anti-CD3 alone (**a**) (TCRζ-ZAP70 lag: *n =* 36 MC; TCRζ-c-Cbl lag: *n =* 39 MC; ZAP70-c-Cbl: *n =* 35 MC) or with anti-CD3 + anti-CD28 (**d**) (TCRζ-ZAP70 lag: *n =* 35 MC; TCRζ-c-Cbl lag: *n =* 35 MC; ZAP70-c-Cbl: *n =* 36 MC) antibodies. 120 time-lapse images were acquired every 3s at 21°C using TIRF microscope. **a** and **d**— Max intensity projection of representative Jurkat T cells stimulated as indicated above. **b** and **e**– Time-lapse montage of representative microclusters from a and d, as indicated by white boxes, respectively. **c** and **f**— Representative relative intensity plot of the individual microclusters indicated in a and d, respectively. **g**-**i**— Kinetic lags measured between proteins as indicated in the figure under either anti-CD3 (>6 cells) or anti-CD3+anti-CD28 (>4 cells) stimulation. Data presented as mean ± SEM. Populations were analyzed using Student’s (TCRζ-ZAP70 lag), and Welch’s t-tests (TCRζ-c-Cbl lag and ZAP70-c-Cbl lag). \**p* < 0.05, \*\**p* < 0.01. Scale bar: 5μm.

### CD28 co-stimulation mediated acceleration of TCRζ-ZAP70 kinetic lag is abolished in CD28 KO cells

To further confirm that CD28 co-stimulation is required for accelerating the recruitment of ZAP70 to TCRζ in the MC, we decided to use CD28 KO primary mouse cells. To investigate the kinetic lag between TCRζ and ZAP70 in the absence of CD28, CD28 KO mouse primary cells were transfected to express TCRζ-Emerald and ZAP70-Apple. Time-lapse imaging on TIRF microscopes and fluorescence intensities calculated from those movies showed the presence of a kinetic lag between recruitment of ZAP70 and TCRζ when the cells were stimulated with anti-TCR antibody alone (Fig. 4a-c). This mirrors the results observed in mouse primary T cells from WT mice. However, the CD28 KO cells failed to show an accelerated recruitment of ZAP70 to TCRζ in the presence of CD28 co-stimulation added to TCR stimulation (Fig. 4d-f). The mean kinetic lag did not show a statistically significant change between TCR stimulation alone or in combination with CD28 so-stimulation conditions and were recorded at ∼35s and ∼41s respectively (Fig. 4g). These results confirm that CD28 is required for the observed acceleration of the kinetic lag between TCRζ and ZAP70.

**Figure 4:**
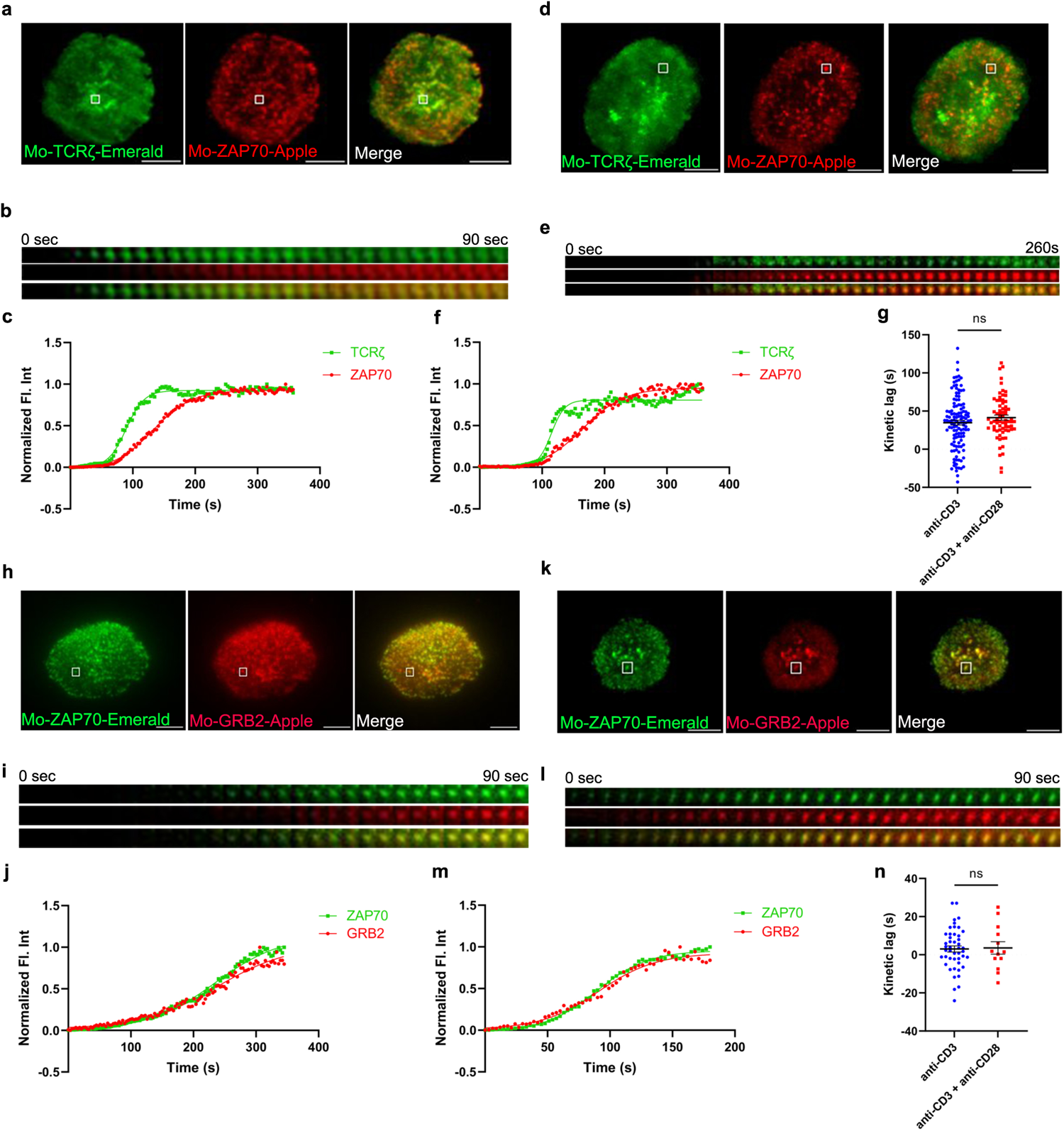
Acceleration of the TCRζ-ZAP70 kinetic lag mediated by CD28 is abolished in CD28 KO cells. Mouse primary T cells from mice genetically manipulated to lack CD28 were transfected to express TCRζ-Emerald (green) and ZAP70-Apple (red) [**a**-**g**] or ZAP70-Emerald (green) and GRB2-Apple (red) [**h**-**n**]. Cells were activated on coverslips coated with either anti-CD3 alone (**a** and **h**) (TCRζ-ZAP70 lag: *n =* 124 MC, >6 cells; ZAP70-GRB2 lag: *n =* 46 MC, >9 cells) or with anti-CD3 + anti-CD28 (**d** and **k**) (TCRζ-ZAP70 lag: *n =* 69 MC, >5 cells; ZAP70-Grab2 lag: *n =* 16 MC, >7 cells) antibodies. 120 time-lapse images were acquired every 3s at 21°C using TIRF microscope. **a, d** and **h, k**— Max intensity projection of representative CD28-deficient primary T cells stimulated as indicated above. **b, e** and **i, l**— Time-lapse montage of representative microclusters from a, d and h, k, respectively, as indicated by white boxes. **c, f** and **j, m**—Representative relative intensity plot of the individual microclusters indicated in a, d and h, k respectively. **g** and **n**— Kinetic lags measured between TCRζ and ZAP70 with indicated stimulatory antibodies. Data presented as mean ± SEM. Populations were analyzed using Welch’s (TCRζ-ZAP70 lag), and Student’s t-tests (ZAP70-GRB2 lag). \**p* < 0.05, \*\**p* < 0.01. Scale bar: 5μm.

We further decided to investigate if CD28 was necessary for the decreased kinetic lag between ZAP70 and GRB2, which was observed in murine WT primary cells and contrasted with our findings in Jurkat T cells. To address this question CD28 KO primary mouse cells were transfected to express ZAP70-Emerald and GRB2-Apple. Time-lapse experiments showed a rather negligible delay between recruitment of ZAP70 and GRB2 with TCR stimulation alone in these CD28 KO cells (Fig. 4h-j) and no significant change was observed in the delay when CD28 KO cells received TCR stimulation in presence of CD28 co-stimulation (Fig. 4k-m). This supports the conclusion that CD28 engagement was responsible for the decrease in the kinetic lag we observed above.

### CD28 regulates Lck localization and activation

As discussed, upon TCR-pMHC ligation Lck phosphorylates the ITAM motifs in the TCRζ chain and creates docking sites for ZAP70. Lck phosphorylates and activates ZAP70 when it is recruited to the cytoplasmic tail of the TCRζ chain^20^. CD28 is known also to interact with the Src family tyrosine kinase, Lck.^14,21^ Because of the upstream role of Lck at the TCR, as well as its interaction with CD28, we decided to test the hypothesis that CD28 could further regulate Lck dynamics. Such an effect could be central to the mechanism of function CD28 on co-stimulation and TCR activation. We stimulated Jurkat T cells expressing Lck-Emerald and ZAP70-Apple with anti-CD3ε antibody in the absence or presence of CD28 co-stimulation and performed qualitative and quantitative analysis of Lck dynamics at the ZAP70 MCs using time-lapse imaging with TIRF microscopy (Fig. 5a). We observed first that compared to TCR stimulation alone, in the presence of CD28 co-stimulation Lck was increasingly enriched in the MCs marked by ZAP70, as indicated by the change in normalized fluorescence intensities of Lck over time (Fig. 5b). This observation along with the evidence of interaction between CD28 and Lck prompted us to ask if CD28 alone is sufficient to recruit Lck and form Lck clusters. To test this question, Jurkat T cells expressing CD28-turquoise, Lck-Apple, and TCRζ-YFP were stimulated with CD3ε or CD3ε and anti-CD28 or anti-CD28 alone. CD3ε stimulation alone showed reduced Lck clusters and reduced recruitment of Lck to TCRζ. TCR stimulation combined with CD28 co-stimulation showed abundant Lck clusters and considerable colocalization with TCRζ. Interestingly, CD28 stimulation alone also led to formation of a considerable amount of Lck clusters (Supp. Fig. 4). These results suggest that CD28 can indeed recruit Lck and form Lck clusters, which can in the presence of anti-CD3ε stimulation enhance TCR-recruited Lck.

**Figure 5:**
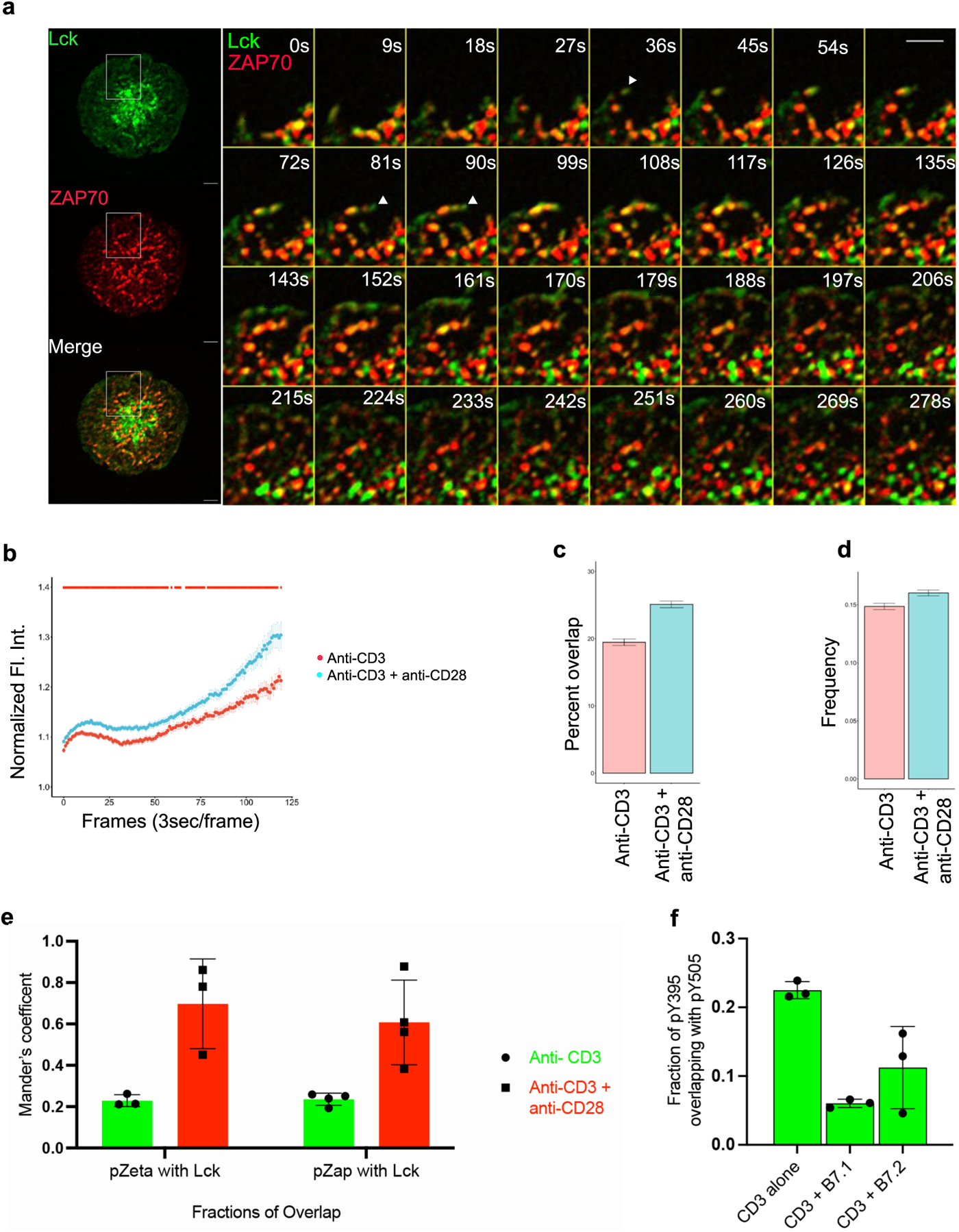
Lck localization and activation is regulated by CD28 co-stimulation. Jurkat T cells were transfected to express Lck-Emerald (green) and ZAP70-Apple (red). Cells were activated on coverslips coated with either anti-CD3 alone or with anti-CD3 + anti-CD28. 120 time-lapse images were acquired every 3s at 21°C using TIRF microscope (a-d). **a**— Max intensity projection of a representative Jurkat T cell becoming activated on coverslip-bound anti-CD3 and anti-CD28 antibody (Left). Time-lapse montage (Right) of a section of the cell as indicated by white box shows early dynamics and priming action of Lck at sites where a ZAP70 microcluster would form. **b**– Normalized fluorescence intensity over time of Lck at ZAP70 microclusters in cells stimulated as indicated. **c**—Percent overlap of Lck at ZAP70 microclusters as a function of time in cells stimulated as indicated. **d**— Frequency of change between “Lck ON” and “Lck OFF” state of ZAP70 microclusters in cells stimulated as indicated. **e**— Jurkat T cells were stimulated with anti-CD3 alone or with anti-CD3 + anti-CD28 coated coverslips for 5 minutes and then stained with total Lck, TCRζ-pY142 and ZAP70-pY319. Graph shows colocalization of Lck with TCRζ-pY142 and ZAP70-pY319 and in cells stimulated as indicated. **f**— Jurkat T cells were stimulated with anti-CD3 alone, anti-CD3 + B7.1 and anti-CD3 + B7.2 coated coverslips for 5 minutes and then stained with Lck-pY395 and Lck-pY505. Graph shows fraction of overlap of Lck-pY394 (activating phosphorylation) with Lck-pY505 (inhibitory phosphorylation) on cells stimulated as indicated. Data presented as mean ± SEM. Populations were analyzed using Welch’s (TCRζ-ZAP70 lag), and Student’s t-tests (ZAP70-GRB2 lag). \**p* < 0.05, \*\**p* < 0.01. Scale bar: 5μm.

Time-lapse montages and movies showed that Lck was present at the leading edge of the MC formation front and with extended dwell time move in the plasma membrane to points where ZAP70 will recruit and form a stable MC (Fig. 5a left panel). Quantification of Lck dwell-time over the course of the time-lapse imaging revealed that Lck showed a statistically significant increase in percent localization overlap with ZAP70 when Jurkat cells were stimulated with a combination of TCR stimulation and CD28 co-stimulation compared to cells receiving TCR stimulation alone (Fig. 5c). Another striking pattern was the “Lck on” and “Lck off” state of ZAP70 MCs as Lck frequently visited and then disappeared from the ZAP70 MC. Additionally, Lck could be seen actively shuttling between ZAP70 clusters (Fig. 5a right panel). Such dynamic oscillations observed for Lck on stable ZAP70 clusters could explain Lck’s role in activating newly recruited ZAP70 in the MC. Therefore, we measured the frequency of change between “Lck off” and “Lck on” state of ZAP70 clusters in cells that stimulated with either TCR stimulation alone or TCR stimulation along with CD28 co-stimulation. Consistent with the localization and dwell-time data, CD28 co-stimulation also led to a significant increase in frequency of change between “Lck on” and “Lck off” state of ZAP70 clusters (Fig. 5d). This suggests that CD28 co-stimulation enhanced the frequency of Lck visits to the ZAP70 MCs and therefore, significantly increased the dwell time of Lck in the MCs.

We next examined if Lck enrichment mediated by CD28 co-stimulation could be correlated with activation of ZAP70 and TCRζ. Colocalization analysis of Lck and active pZAP70 Y319 and pTCRζ Y142 showed a significant increase in overlap of Lck and active ZAP70 and TCRζ when cells were stimulated with anti-CD28 along with anti-CD3ε compared to that of anti-CD3ε stimulation alone (Fig. 5e). It is also relevant to ask whether CD28 co-stimulation could regulate Lck activation in addition to Lck localization. Lck activity is tightly regulated by two key phosphorylations. Lck undergoes an activation phosphorylation at tyrosine 394 in the kinase domain and a dephosphorylation at tyrosine 505 residue in the inhibitory region to acquire a fully open and active conformation. However, the singly phosphorylated (pY505) alone or doubly phosphorylated forms (pY394 and pY505) of Lck remain in a closed and inactive conformation ^22–24^. Our analysis revealed that a combination of TCR stimulation and CD28 co-stimulation resulted in a significantly diminished overlap of pY394 with pY505 compared to TCR stimulation alone (Fig. 5f). These results indicate that CD28 co-stimulation shifts the total Lck pool towards the active form of pY394 Lck.

### CD28 co-stimulation results in increase in ZAP70 phosphorylation and results in earlier calcium flux

CD28 co-stimulation is known regulate several signaling pathways which are initiated at its cytoplasmic tail^13,25^. These pathways regulate several signaling outcomes such as activation of downstream proteins, Ca^2+^ flux, actin remodeling, cytokine production, proliferation, and transcription^10^. We specifically focused on early events downstream of TCR engagement that coincide and correlate with accelerated ZAP70 recruitment such as TCRζ phosphorylation, ZAP70 activation, PLCγ1 activation and Erk phosphorylation. Activation of these signaling proteins were analyzed as early as 15s and up to 300s of activation with either TCR stimulation alone or in combination with CD28 co-stimulation. We did not see any change in phosphorylation of TCRζ, which is not surprising because it is upstream of ZAP70 recruitment (Fig. 6a and b). Erk activation also remained unchanged suggesting that the Erk pathway may not be affected by accelerated ZAP70 recruitment (Fig. 6a and e). Consistent with our kinetic lag data, ZAP70 showed a robust increase in phosphorylation in the presence of CD28 co-stimulation compared to TCR stimulation alone. This increase was significant as early as 30s after activation and remained consistently elevated through 300s after activation. (Fig. 6a and c). These data confirmed a clear role for CD28 co-stimulation in early activation of ZAP70. In contrast to our expectation, we did not see a change in phosphorylation and activation of PLCγ1 (Fig. 6a and d). Since the ratio of PLCγ1 that is recruited to the MC and becomes activated is relatively quite low in comparison to the total cellular PLCγ1, we thought that it was possible that CD28-mediated increase in PLCγ1 activation may be difficult to detect by Western blot. Therefore, we decided to investigate whether Ca^2+^ flux, which is downstream of PLCγ1 activation, is altered in the presence of CD28 co-stimulation. Using GCaMP6s as a genetically encoded calcium reporter, we investigated the kinetic lag between the recruitment of TCRζ and initiation of Ca^2+^ flux upon TCR stimulation either in the presence or absence of CD28 co-stimulation. Our results definitively showed a significant acceleration in the onset of Ca^2+^ flux in the presence of CD28 co-stimulation (Fig. 6f). These results suggest that one the of the functional consequence of accelerated ZAP70 recruitment and ZAP70 activation could be an earlier onset of Ca^2+^ flux, which may optimize TCR stimulation

**Figure 6:**
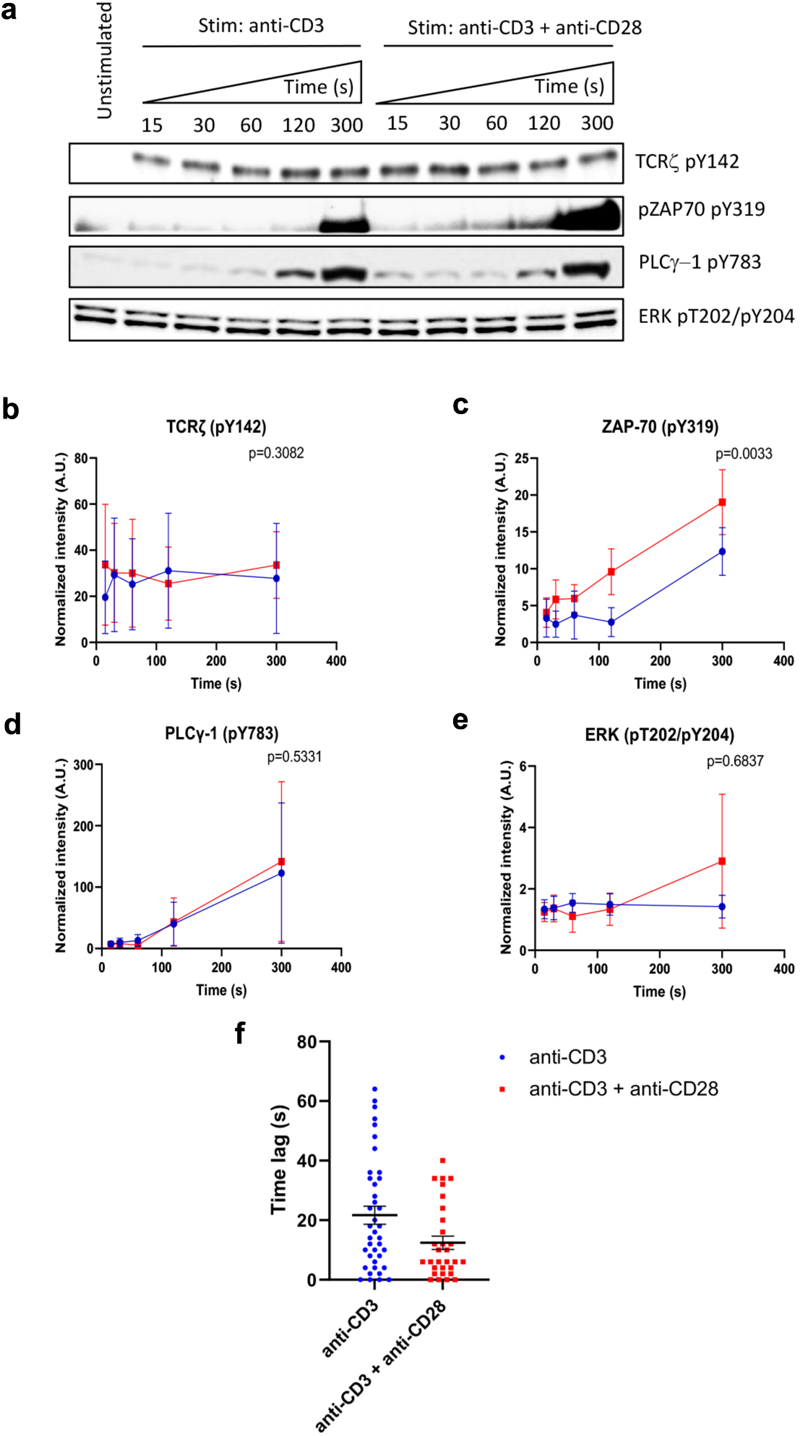
CD28 co-stimulation increases ZAP70 activation and accelerates onset of calcium flux. **a-e**— Jurkat cells were stimulated on plate bound anti-CD3 alone or anti-CD3 + anti-CD28 for indicated durations. Lysates were resolved by SDS-PAGE and Western blot was performed with indicated antibodies (**a**). **b-e**— Band intensities were calculated using ImageJ and normalized intensities were plotted for indicated proteins at each time point. **f**— Jurkat T cells were transfected to express the calcium indicator GCaMP6s (green) and TCRζ-Apple (red). Cells were activated on coverslips coated with either anti-CD3 alone or with anti-CD3 + anti-CD28. 200 time-lapse images were acquired every 2s at 21°C using TIRF microscope. Time lag between onset of TCRζ and GCaMP6s intensities in cells stimulated as indicated. Data presented as mean ± SEM (*n =* 3 for all experiments). Populations analyzed using 2-way ANOVA (**b**-**d**) and Mann-Whitney test (**f**). \**p* < 0.05.

## Discussion

The CD28 family of receptors in T cells include co-stimulatory molecules such as CD28 and ICOS (inducible co-stimulator), and co-inhibitory molecules such as CTLA-4 and PD1. Signaling from these co-stimulatory and co-inhibitory molecules integrate with TCR signaling to tune an optimal T cell response. An extensive amount of research has uncovered the role of CD28 co-stimulation in regulating immune response, ligand discrimination, T cell fate and T cell development^10,13^. The binding of the CD28 ligands, B7.1 and B7.2 expressed on APCs, to the extracellular domain of CD28 leads to signaling events from proteins bound to its evolutionary conserved cytoplasmic tail. Two major signaling motifs in the cytoplasmic tail of CD28 include the membrane proximal YMNM domain and the membrane distal proline-rich PYAP domain. Phosphorylation of the tyrosine in the YMNM motif by either of the Src family kinases, Lck or Fyn creates a docking site for SH2 domain-containing proteins such as GRB2, GADS and the p85 subunit of PI3-kinase. Binding events at the PYAP domain include recruitment and activation of Lck and proteins interacting at this site also include Filamin A, GRB2 and GADS via their SH3 domains. There is thus some functional redundancy of molecular binding to these signaling motifs in CD28. Despite our elaborate understanding of these molecular interactions at CD28 and their functional relevance, a definitive mechanism of function for CD28 is not clear.

TCR-mediated signaling initiates formation of sub-micron sized molecular aggregates called microclusters (MC). These molecular structures, now known to be phase-separated condensates, contain transmembrane receptors, co-receptors, membrane-associated and cytoplasmic kinases, enzymes and adapter proteins^2,4,26–29^. Complex phosphorylation and dephosphorylation events, and the spatial arrangements and stoichiometry of these molecules regulate signaling from these MCs from the initiation of activation. These MCs are functional signaling elements which over time coalesce to form the immunological synapse and propagate downstream cellular responses. Using super-resolution microscopy, we have previously shown that the MC contains two major domains which are spatially and functionally distinguishable. The receptor domain is composed of the TCRζ and the kinase, ZAP70, whereas the signaling domain contains adapter and signaling proteins such as LAT, GADS, SLP76, ADAP and PLCγ1^3^. We further elucidated the spatio-temporal relationship, stepwise recruitment, and time delay between recruitment of these receptor and signaling domain proteins in the MC^3^.

In our current study we found that CD28 co-stimulation specifically accelerated the recruitment of ZAP70 to TCRζ by decreasing the kinetic lag between these two molecules in both Jurkat and murine peripheral T cells. The kinetic lag, which is defined as the time between two molecules reaching their half-maximum fluorescence intensities, can be used as a surrogate for mapping the sequential recruitment of fluorophore-tagged proteins. CD28 co-stimulation increased the overall percentage of MCs that had shorter TCRζ-ZAP70 kinetic lags compared to MCs which had longer kinetic lags. This skewing of MCs towards a shorter kinetic lag between the receptor and the kinase supported our hypothesis that CD28 co-stimulation predominantly accelerates recruitment of ZAP70 to the receptor in most MCs. Intriguingly, the MCs in the short kinetic lag bin (0sec to 25s) which received CD28 co-stimulation along with CD3 stimulation as opposed to TCR stimulation alone also started to form earlier during cell spreading on the stimulatory surface. We propose that these alterations in ZAP70 recruitment during the formation of MCs underlie the substantial changes in activation of ZAP70 and relay of cellular signals in the presence of the co-stimulatory signal. The most upstream effect of CD28 engagement appears to be the shortened kinetic lag between TCRζ and ZAP70, which has been consistently observed in Jurkat T cells and murine primary T cells.

The kinetic lag between ZAP70 and other signaling domain proteins such as GRB2 and SLP76 in Jurkat cells did not show a significant change in response to CD28 co-stimulation. However, in murine T cells CD28 co-stimulation also significantly decreased the lag between ZAP70 and GRB2. On a molecular basis this divergent observation might be attributed to loss-of-function mutations, such as PTEN and CTLA4, in T cell receptor pathway genes in Jurkat T cells^30^. These genetic differences might affect the membrane abundance of CD28 as a result of PTEN loss, and/or might dampen CD28 signaling by CTLA4-mediated sequestration of a PP2A, a negative regulator of CD28 signaling^30–35^. Despite the absence of these above-mentioned signaling pathways in Jurkat cells, the shortened kinetic lag between TCRζ and ZAP70 in both Jurkat and normal murine T cells showed significant downstream effects, such as accelerated Ca^2+^ flux and ZAP70 activation. These results suggest that the kinetic lag between TCRζ and ZAP70 could be the most important step downstream of CD28 co-stimulation.

Our findings also reveal a molecular mechanism, direct regulation of Lck recruitment, activation, and dynamics at the MC, which explains how accelerated ZAP70 recruitment and activation are achieved through CD28 co-stimulation. Our results showed that Lck recruitment and enrichment in the MC, and activation of Lck was markedly increased upon CD28 co-stimulation. This conclusion depends on our analysis of Lck phosphorylation on two Lck residues, Y394 and Y505. It has been found that Y394 auto-phosphorylation leads to Lck activation. Phosphorylation at Y505 results in a closed conformation of Lck, thus rendering Y394 inaccessible for auto-phosphorylation^23^ Lck molecules can be found in three different states (a) open and active (unphosphorylated at Y505 and phosphorylated at Y394), (b) open and primed (unphosphorylated at Y505 and unphosphorylated at Y394), and (c) closed and inactive (phosphorylated at Y505 and unphosphorylated at Y394)^36,37^. Our analysis of phosphorylation states of Lck revealed a skewed distribution among the above-mentioned forms upon CD28 co-stimulation, and favored an increase in the amount of Lck present in the open and active conformation. As described above, the open and active form of Lck will exist as phosphorylated at Y394 and unphosphorylated at Y505. Our results also demonstrate a decrease in the doubly phosphorylated (p-Y394/p-Y505) species of Lck. **T**his suggests that CD28 co-stimulated cells may have the phosphorylation equilibrium shifted towards singly phosphorylated Y394, which is most like to be the fully active form of Lck. We propose that CD28-mediated clustering of the activated form of Lck could be a proximal and important mechanism by which CD28 co-stimulation enhances the efficiency of TCR signaling.

For these proximal molecular events triggered by CD28 co-stimulation to shape TCR response, a sustained activation of downstream signaling molecules ensures that cellular responses will proceed. We characterized the downstream consequences of accelerated recruitment of ZAP70. We observed an early activation of ZAP70, and we observed an impressive early onset of Ca^2+^ flux, as a cellular response, upon CD28 co-stimulation compared to TCR stimulation alone. However, we did not observe any significant increase in Erk phosphorylation or IL2 production (data not shown). These results do not necessarily reflect a limitation in propagation of the effect of CD28 co-stimulation at the scale of cellular response. Our experimental system used anti-TCR antibody, which results in a strong TCR activation without much ability to tune the affinity and strength of the signal. To truly appreciate the effect of CD28 co-stimulation in shaping TCR signaling, we would need to use a variable affinity antigen-based TCR activation system, which was beyond the scope of this study.

Our results suggest a mechanism of CD28 co-stimulation that can be better explained by means of the Kinetic Proof-Reading model of T cell activation. TCR signaling can be thought to be comprised of several intermediate and contingent steps which translate the TCR-pMHC (receptor-ligand) interaction into cellular response such as Ca^2+^ flux, gene expression, and cytokine production. CD28 co-stimulation may increase the efficiency of one or more of those intermediate steps to allow the signaling cascade to propagate efficiently without halting or reversing, thereby allowing the TCR to respond to low affinity antigens, allow self vs. non-self-antigen recognition, and allow limited duration ligand-receptor interaction to trigger signaling. The mechanism of function of CD28 revealed in this study defines a molecular mechanism that defines how CD28 signaling can tune TCR signaling and allow for sustained signaling resulting in cellular response.

## Methods

### DNA constructs

Human expression constructs: TCRζ-Emerald, ZAP70-Apple, ZAP70-Emerald, GRB2-Emerald, SLP76-Emerald, GADS-Emerald, and c-Cbl-YFP constructs were described previously^3,28,38^. pGP-CMV-GCaMP6s (GCaMP6) construct was a gift from Douglas Kim (Addgene plasmid # 40753). TCRζ-Turquoise was generated by cloning turquoise sequence in plasmid expressing TCRζ with AgeI-NotI sites. TCRζ-YFP was generated by cloning TCRζ cDNA sequence from TCRζ-Emerald vector into mYFPN1 plasmid (Clontech) vector using NheI-AgeI sites. TCRζ-Apple was generated by cloning Apple sequence into a plasmid expressing TCRζ using AgeI-NotI sites. Lck-Apple and Lck-Emerald constructs were generated by cloning AgeI-NotI digested Apple and Emerald sequence, respectively, into a plasmid containing the Lck cDNA sequence. CD28-Apple and CD28-turquoise constructs were generated by cloning AgeI-NotI-sdigested Apple and Turquoise sequences respectively into a plasmid containing CD28 cDNA sequence. Murine expression constructs: Mouse TCRζ-Emerald construct was generated by cloning mouse-TCRζ cDNA sequence into an Emerald-expressing plasmid using AgeI-NheI sites. Mouse ZAP70-Emerald and ZAP70-Apple constructs were generated by cloning mouse-ZAP70 cDNA sequence into an Emerald or Apple expressing plasmids, respectively, using AgeI-NheI sites. Mouse GRB2-Apple construct was generated by cloning mouse-GRB2 cDNA sequence into an Apple expressing plasmid using AgeI-NheI sites.

### Reagents

Human anti-CD3ε (Clone: HIT3a), human anti-CD3ε (Clone: UCHT1), and human anti-CD28 (Clone: CD28.2) monoclonal antibodies were customized for concentration and purchased from BD Pharmingen and were used to coat glass bottom chambers for imaging assays and phosphorylation assays in Jurkat T cells. Human anti-CD43 (Clone: S11) was purchased from BioLegend (Cat. No.143201) and was used to coat glass bottom chambers for imaging studies. Mouse anti-CD3ε (Clone: 145-2C11, Cat. No. 553058) and mouse anti-CD28 (Clone: 37.51, Cat. No. 553294) monoclonal antibodies were purchased from BD Pharmingen and were used to coat glass bottom chambers for imaging assays in murine T cells. Natural ligands for CD28 used in this study are B7.1 (Cat. No. 310-32-100UG) and B7.2 (Cat. No. 310-33-100UG) from Preprotech. The following antibodies were used for immunostaining: anti-Lck (Clone: 3A5, Millipore), anti-Lck pY505 (BD Transduction, Cat. No. 612390)), anti-LckpY394 (Cell signaling, Cat. No. 2101S), anti-ZAP70pY319 (Cell Signaling, Cat. No. 2717S), anti-p-TCRζY142 (BD Pharmingen, Cat no. 558402). Alexa Fluor secondary antibodies and secondary antibodies used in Western blot assays were purchased from Thermo Fisher Scientific. Imaging buffer (20 mM Hepes pH 7.2, 137 mM NaCl, 5 mM KCl, 0.7 mM Na2HPO4, 6 mM D-glucose, 2 mM MgCl2, 2 mM CaCl2, 1% BSA) has been described before^3^.

### Phosphorylation and Western blotting assays

Jurkat T cells were activated on 96-well plates (coated with 2ug/ml HIT3a antibody with or without 2ug/ml anti-CD28 antibody in 1X PBS) in imaging buffer (described above) at 500,000 cells per well. Prior to stimulation, 30e6 cells were washed twice with 1X PBS before being resuspended at a concentration of 10e6 cells/ml in RPMI without supplements and rested on ice for 15 minutes. Cells were then resuspended in imaging buffer at 10e10 cells per ml and were used for stimulation. Pre-heated lysis buffer (20mM Tris-HCl (pH8), 2mM EDTA, 2mM Na3VO4, 20mM DTT, 2% SDS, and 20% glycerol in ultra-pure water) was added at specified times after stimulation. Lysates were then transferred to eppendorf tubes and heated at 100℃ for 10 minutes, sonicated, and stored at −20 °C. Protein lysates were separated by SDS-PAGE. Anti-TCRζ (Santa Cruz Biotechnology, Cat. No. SC-1239), anti-ZAP70 (Cell Signaling, Cat. No. 2705S), anti-PLCγ-1(Cell Signaling, Cat. No. 2822S), anti-Erk (Cell Signaling, Cat. No. 4685S), anti-p-TCRζ-Y142 (BD Pharmingen, Cat no. 558402, anti-p-ZAP70-Y319 (Cell Signaling, Cat. No. 2717S), anti-p-PLCγ-1-Y783 (Cell Signaling, Cat. No. 2821S) and anti-p-Erk-T202/Y205 (Cell Signaling, Cat. No. 43702) were used to detect total and phosphorylated forms of TCRζ, ZAP70, PLC-γ-1 and Erk respectively. HRP-conjugated secondary antibodies and Super-Signal West chemiluminescent substrates (Thermo Scientific) were used for detection. Western blot band Intensities were measured using ImageJ. Band intensities of phosphorylated protein bands were normalized to the intensity of total protein bands, and further normalized to unstimulated controls.

### Cell culture, transfection and preparation of Jurkat cells

Culture and maintenance of Jurkat E6.1 cells have been described previously^17^. E6.1 Jurkat cells were cultured in RPMI (11875–093; Life Technologies) supplemented with10% fetal bovine serum (26140–079; Life Technologies). For transient transfection 1e6 Jurkat cells were transfected with 2 μg DNA using the LONZA electroporator (program X-001) and Nucleofector Kit V (Lonza, catalog no. VCA-1003) 24 h prior to imaging. Before imaging, Jurkat cells were spun down and resuspended in imaging buffer.

### Cell culture, transfection and preparation of primary mouse T cells

Pan CD4^+^ cells were isolated from spleens from gender and age matched C57BL/6 and CD28 KO (B6.129S2-Cd28tm1Mak/J, Strain #: 002666) mice from Jackson laboratories using Pan-T Cell Isolation Kit (130–095–130; Miltenyi Biosciences). Lymphocytes were cultured in RPMI supplemented with 10% FBS and 50 μM beta-mercaptoethanol (RPMI media) Lymphoblasts were generated by activation with plate bound murine anti-CD3ε antibody (BD Biosciences, Clone: 145-2C11) in the presence of 100 U/ml of human IL-2 and soluble anti-CD28 antibody (BD Biosciences, Clone: 37.51). T lymphoblasts were cultured at 37 °C in 5% CO2 for 3 days in exponential growth phase. Cells were then cultured for an additional day in RPMI media and in the presence of 100 U/ml of IL-2 only. Cells were then rested in RPMI media without IL2 and CD28 for an additional day before transfection. For electroporation, 5×1e6 cells were transfected with 5μg/plasmid DNA/million cells using a LONZA electroporator (program X-001) and LONZA electroporation kit for primary mouse T cells (VPA-1006; LONZA) according to the manufacturer’s protocol. Cells were then incubated in RPMI media for 8-12 hrs. Before transfection cells were spun down and resuspended in imaging buffer (described above).

### Live cell imaging

Antibody-coated chambers were prepared as described previously^27^. Jurkat cells were added to the imaging buffer in the antibody(ies) and/or ligand coated 8-well coverslip chambers (Lab-Tek, Thermo Fisher) and imaged at room temperature (21 °C). For coating coverslips antibodies or natural ligands were used at 10ug/ml for kinetic assays or at 5ug/ml for Ca^2+^ flux assays in 1X-PBS. TIRF images from live cells were collected with a Nikon Ti-E inverted microscope, using a 100X SR Apochromat TIRF objective lens (1.49 numerical aperture), and an Andor iXon Ultra 897 EM charge-coupled device camera (512 X 512 pixels, 16 μm pixel). Time-lapse images were collected at 3 s/frame. For Ca^2+^ imaging assays, TIRF images from live cells were collected with a Nikon Ti-E inverted microscope, using a 60X SR Apochromat TIRF objective lens (1.49 numerical aperture), and an Andor iXon Ultra 897 EM charge-coupled device camera (512 X 512 pixels, 16 μm pixel). Time-lapse images were collected at 2 s/frame.

### Immunofluorescence imaging

Jurkat T cells were stimulated by addition in imaging buffer to the antibody/ies and/or ligand-coated 8-well coverslip chambers (Lab-Tek, Thermo Fisher). Incubation was for 5 minutes at room temperature (21 °C). For coating coverslips, antibodies or natural ligands were used at 10ug/ml in 1X-PBS. Cells were then fixed, permeabilized and incubated with blocking solution [1X PBS with 10% FBS (Sigma-Aldrich) and 0.01% sodium azide (Sigma-Aldrich)] for immunofluorescence staining as described previously^3^. Cells were then washed 3 times with 1X PBS and then incubated with primary antibodies in blocking solution followed by 3 washes with 1X PBS and incubation with secondary antibodies prepared in blocking solution. For experiments in Fig.3e, Alexa 488, Alexa 568, and Alexa 647 were used to detect pZetaY142, Lck and pZAP70Y319, respectively. For experiments in Fig.3f, Alexa 488 and Alexa 546 were used to detect pLckY394 and pLckY505, respectively. Cells were finally washed with 1X PBS for 3 times and kept immersed in 1XPBS while being imaged by TIRF microscopy using a 100X SR Apochromat TIRF objective lens (1.49 numerical aperture), and an Andor iXon Ultra 897 EM charge-coupled device camera (512 X 512 pixels, 16 μm pixel).

### Image analysis

Image J was used for generating the kymographs and maximum projection images from time-lapse movies from live cell experiments and 2D images from immunofluorescence assays which were acquired on the TIRF microscope. Kinetic analysis of individual microclusters, normalization of raw intensity values and sigmoidal non-linear curve fitting has been previously described^3^. For co-localization and percent overlap studies Mander’s coefficient was calculated using Image J.

Analysis of ZAP70-labeled TCR MCs from TIRF images was performed using Imaris Bitplane (Oxford Instruments). ZAP70-Apple expressing microclusters were segmented using the Spots module and tracked over time. The normalized Lck TCR enrichment was calculated by quantifying Lck intensity at segmented ZAP70 microclusters normalized to the total cellular Lck intensity.

Lck dwell time was calculated by defining Lck overlap at ZAP70 microcluster as a normalized Lck intensity equal or more than 1.25. The total amount of positive Lck / ZAP70 overlap time was normalized to the total ZAP70 track time to calculate the Lck dwell time on a per-MC track basis. The frequency of change between Lck ON and OFF states at ZAP70 microclusters was calculated by quantifying the number of transitions from Lck / ZAP70 overlap to non-overlap states normalized to the total Zap70 track time.

For analyzing the time lapse imaging data from Ca^2+^ flux assays, ImageJ was used to draw an ROI around each cell for further analysis. ROIs were then analyzed using an in-house custom built MATLAB program which measured fluorescence intensities for individual channels, performed background subtraction, performed data normalization, and then measured time lag between the start of calcium flux and microcluster formation.

## Author Contributions

K.R., R.R., and A.M. performed experiments; K.R., J.Y., and L.E.S. designed the study; K.R., R.R., A.M., A.T., and N.A. performed data analysis; S.P. wrote MatLab programs to analyze data; K.M.M. and J.A.G. generated expression plasmids; K.R. prepared the figures with the help from R.R.; K.R. and L.E.S wrote the manuscript with help from R.R. and A.T.

## Acknowledgements

This research was supported by the Intramural Research Program of the NIH, NCI, CCR. We thank Xufeng Wu and NHLBI for access to their microscope core facility. We thank Paul Randazzo for input on data analysis. We thank Wenmei Li for help with supplying reagents and animals for the study. We thank the NCI CCR Genomics core for sequencing our constructs.

## Supplementary Figures and Legends

**Supplementary Figure 1:**
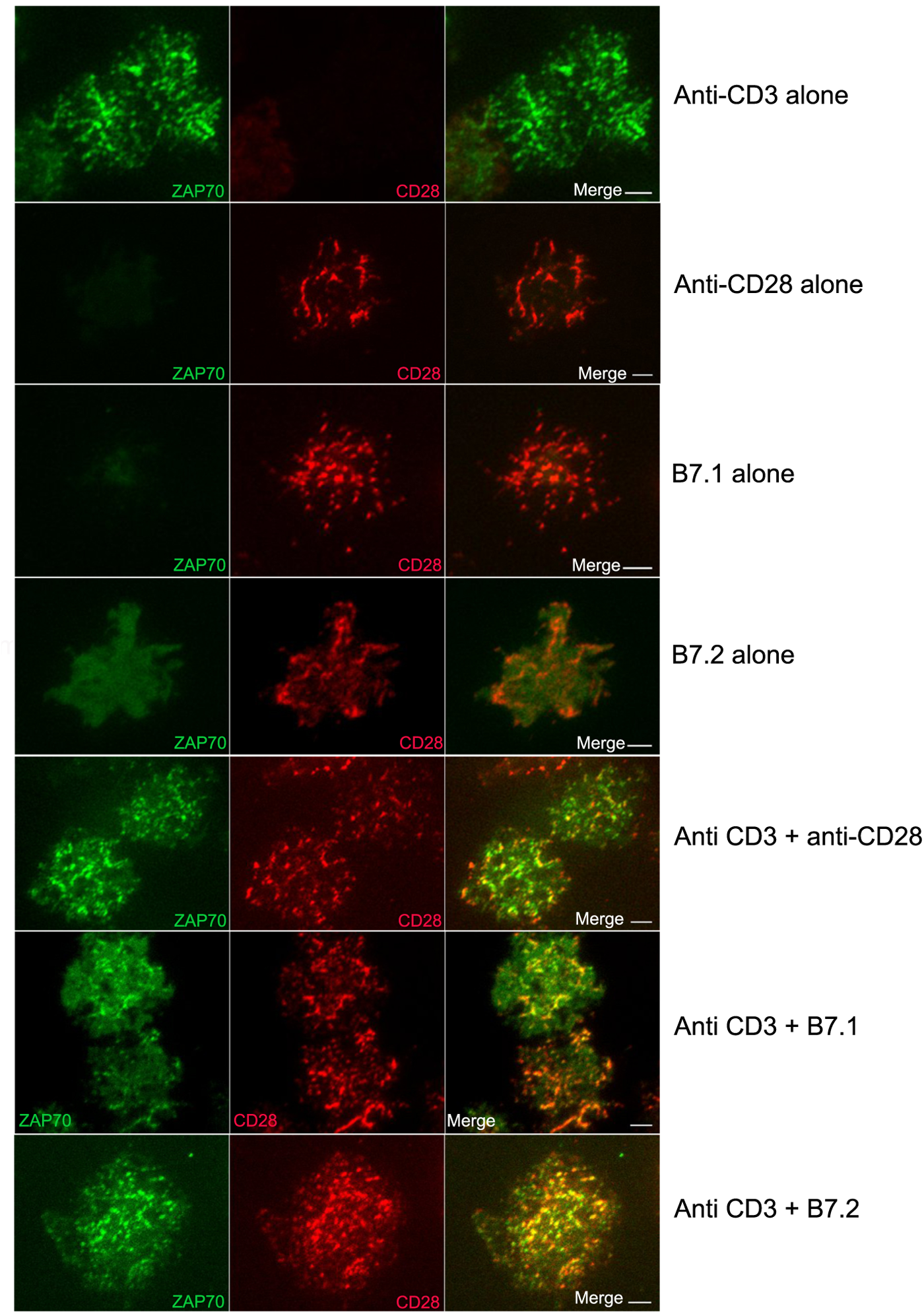
Anti-CD28 antibody and natural ligands for CD28 (B7.1 and B7.2) specifically form CD28 microclusters which colocalize with ZAP70. Jurkat T cells were transfected to express ZAP70-Emerald (green) and CD28-Apple (red). Cells were activated on coverslips coated with indicated antibodies, natural ligands, or combinations. Images were acquired at 21°C using TIRF microscope. Scale bar: 5μm.

**Supplementary Figure 2:**
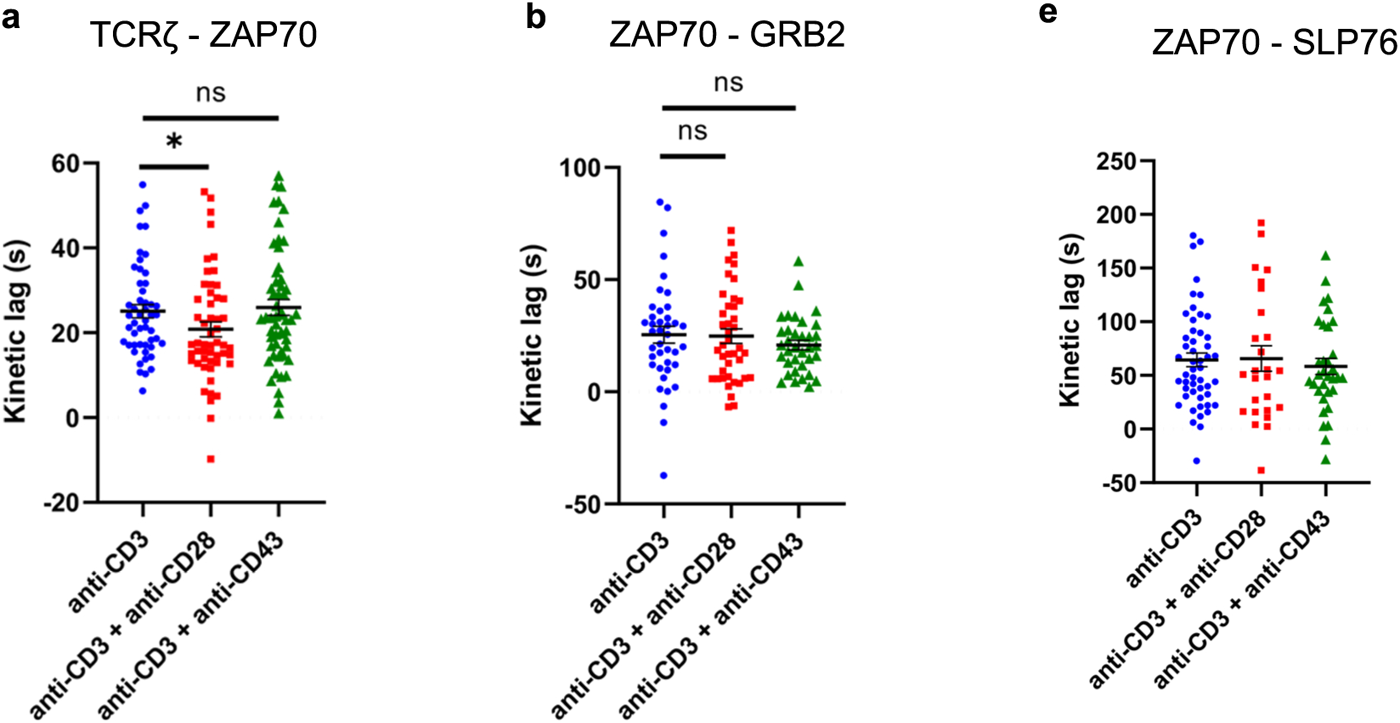
Accelerated recruitment of ZAP70 to TCRζ in the TCR microclusters is specifically driven by anti-CD28 antibody. Jurkat T cells were transfected to express TCRζ-Emerald (green) and ZAP70-Apple (red) or ZAP70-Apple (red) and GRB2-Emerald (green) or ZAP70-Apple (red) and SLP76-Emerald (green) and were activated on coverslips coated with either anti-CD3 alone (TCRζ-ZAP70 lag: *n =* 50 MC, >7 cells; ZAP70-Grb2 lag: *n =* 40 MC, >8 cells; ZAP70-SLP76 lag: *n =* 50 MC, >9 cells), or with anti-CD3 + anti-CD28 (TCRζ-ZAP70 lag: *n =* 52 MC, >7 cells; ZAP70-GRB2 lag: *n =* 41 MC, >11 cells; ZAP70-SLP76 lag: *n =* 26 MC, >6 cells), or with anti-CD3 + anti-CD43 (TCRζ-ZAP70 lag: *n =* 55 MC, >7 cells; ZAP70-GRB2 lag: *n =* 35 MC, >5 cells; ZAP70-SLP76 lag: *n =* 33 MC, >5 cells). 120 time-lapse images were acquired every 3s at 21°C using TIRF microscope. **a-c**— Kinetic lags measured between TCRζ and ZAP70, ZAP70 and GRB2 and ZAP70 and SLP76, respectively with indicated stimulatory antibodies. Data presented as mean ± SEM. Populations were analyzed using Mann-Whitney tests (TCRζ-ZAP70 lag), Welch’s t-tests (ZAP70-GRB2 lag) and Student’s t-tests (ZAP70-SLP76 lag). \**p* < 0.05, \*\**p* < 0.01.

**Supplementary Figure 3:**
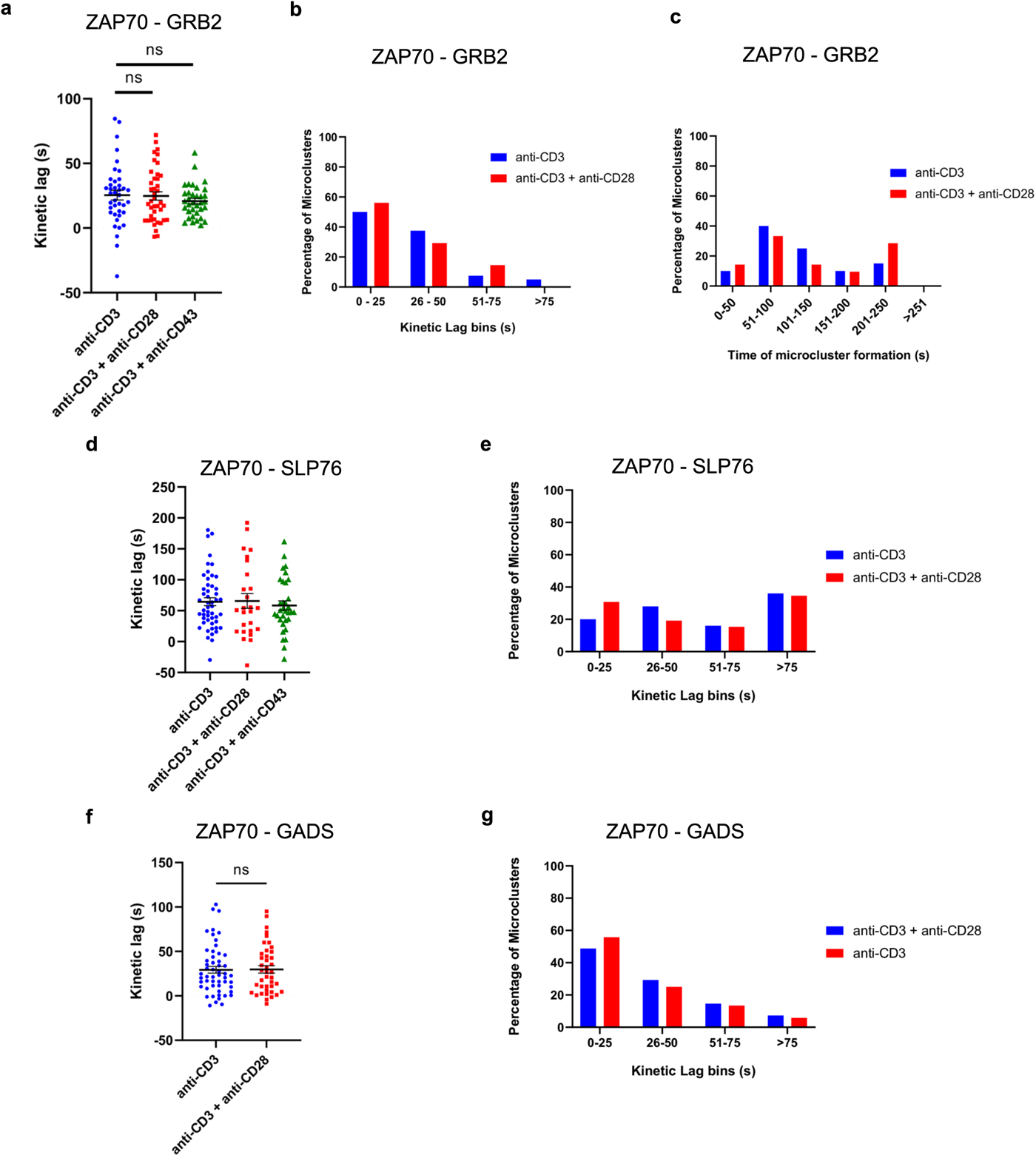
Effect of CD28 co-stimulation is less pronounced in regulating kinetic lag between ZAP70 and signaling domain proteins than kinetic lag between TCRζ and ZAP70. Jurkat T cells were transfected to express ZAP70-Apple (red) and GRB2-Emerald (green) [**a**-**c**], or ZAP70-Apple (red) and SLP76-Emerald (green) [**d**-**e**], or ZAP70-Apple (red) and GADS-Emerald (green) [**f**-**g**] and were activated on coverslips coated with either anti-CD3 alone (ZAP70-GRB2 lag: *n =* 40 MC, >8 cells; ZAP70-SLP76 lag: *n =* 50 MC, >9 cells; ZAP70-GADS lag: *n =* 53 MC, >11 cells), or with anti-CD3 + anti-CD28 (ZAP70-GRB2 lag: *n =* 41 MC, >11 cells; ZAP70-SLP76 lag: *n =* 26 MC, >6 cells; ZAP70-GADS lag: *n =* 41 MC, >9 cells), or with anti-CD3 + anti-CD43 (ZAP70-GRB2 lag: *n =* 35 MC, >5 cells; ZAP70-SLP76 lag: *n =* 33 MC, >5 cells). 120 time-lapse images were acquired every 3s at 21°C using TIRF microscope. **a, d,** and **f**—Kinetic lags measured between ZAP70-GRB2, ZAP70-SLP76, and ZAP70-GADS with indicated stimulatory antibodies. **b, e,** and **g**— Distribution of average kinetic lags across specified time bins for indicated proteins. **c**— Distribution of percentage of microclusters formed at specified time intervals of the first kinetic lag bin (0-25s) in **b**. Data presented as mean ± SEM. Populations were analyzed using Welch’s t-tests (ZAP70-GRB2 lag) and Student’s t-tests (ZAP70-SLP76 lag), Mann-Whitney test (ZAP70-GADS lag). \**p* < 0.05, \*\**p* < 0.01.

**Supplementary Figure 4:**
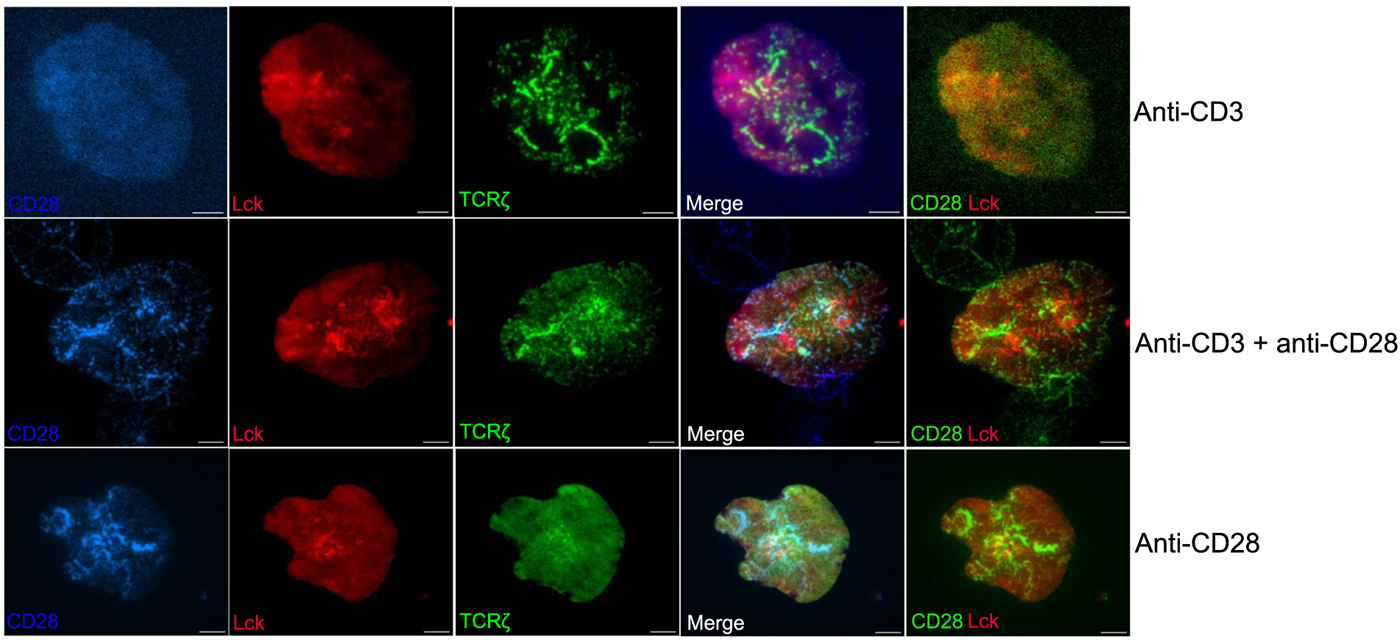
Recruitment of CD28 alone can lead to formation of Lck clusters that colocalize with TCRζ. Jurkat T cells were transfected to express CD28-Turquoise (blue), Lck-Apple (red) and TCRζ-YFP (green) and were activated on coverslips coated with either anti-CD3 alone, or with anti-CD3 + anti-CD28, or with anti-CD28 alone. 120 time-lapse images were acquired every 3s at 21°C using TIRF microscope. Scale bar: 5μm.

## References

1 Balagopalan, L., Kortum, R. L., Coussens, N. P., Barr, V. A. & Samelson, L. E. The linker for activation of T cells (LAT) signaling hub: from signaling complexes to microclusters. J Biol Chem 290, 26422–26429 (2015). 10.1074/jbc.R115.665869

2 Balagopalan, L., Raychaudhuri, K. & Samelson, L. E. Microclusters as T Cell Signaling Hubs: Structure, Kinetics, and Regulation. Front Cell Dev Biol 8, 608530 (2020). 10.3389/fcell.2020.608530

3 Yi, J., Balagopalan, L., Nguyen, T., McIntire, K. M. & Samelson, L. E. TCR microclusters form spatially segregated domains and sequentially assemble in calcium-dependent kinetic steps. Nat Commun 10, 277 (2019). 10.1038/s41467-018-08064-2

4 Su, X. et al. Phase separation of signaling molecules promotes T cell receptor signal transduction. Science 352, 595–599 (2016). 10.1126/science.aad9964

5 Houtman, J. C. et al. Oligomerization of signaling complexes by the multipoint binding of GRB2 to both LAT and SOS1. Nat Struct Mol Biol 13, 798–805 (2006). 10.1038/nsmb1133

6 Grakoui, A. et al. The immunological synapse: a molecular machine controlling T cell activation. Science 285, 221–227 (1999). 10.1126/science.285.5425.221

7 Jenkins, M. K., Ashwell, J. D. & Schwartz, R. H. Allogeneic non-T spleen cells restore the responsiveness of normal T cell clones stimulated with antigen and chemically modified antigen-presenting cells. J Immunol 140, 3324–3330 (1988).

8 Mueller, D. L., Jenkins, M. K. & Schwartz, R. H. An accessory cell-derived costimulatory signal acts independently of protein kinase C activation to allow T cell proliferation and prevent the induction of unresponsiveness. J Immunol 142, 2617–2628 (1989).

9 Gray Parkin, K., et al. Expression of CD28 by bone marrow stromal cells and its involvement in B lymphopoiesis. J Immunol 169, 2292–2302 (2002). 10.4049/jimmunol.169.5.2292

10 Esensten, J. H., Helou, Y. A., Chopra, G., Weiss, A. & Bluestone, J. A. CD28 Costimulation: From Mechanism to Therapy. Immunity 44, 973–988 (2016). 10.1016/j.immuni.2016.04.020

11 Sanchez-Lockhart, M., Kim, M. & Miller, J. Cutting edge: A role for inside-out signaling in TCR regulation of CD28 ligand binding. J Immunol 187, 5515–5519 (2011). 10.4049/jimmunol.1102497

12 Viola, A. & Lanzavecchia, A. T cell activation determined by T cell receptor number and tunable thresholds. Science 273, 104–106 (1996). 10.1126/science.273.5271.104

13 Boomer, J. S. & Green, J. M. An enigmatic tail of CD28 signaling. Cold Spring Harb Perspect Biol 2, a002436 (2010). 10.1101/cshperspect.a002436

14 Dobbins, J. et al. Binding of the cytoplasmic domain of CD28 to the plasma membrane inhibits Lck recruitment and signaling. Sci Signal 9, ra75 (2016). 10.1126/scisignal.aaf0626

15 McKeithan, T. W. Kinetic proofreading in T-cell receptor signal transduction. Proc Natl Acad Sci U S A 92, 5042–5046 (1995). 10.1073/pnas.92.11.5042

16 Rabinowitz, J. D., Beeson, C., Lyons, D. S., Davis, M. M. & McConnell, H. M. Kinetic discrimination in T-cell activation. Proc Natl Acad Sci U S A 93, 1401–1405 (1996). 10.1073/pnas.93.4.1401

17 Balagopalan, L. et al. c-Cbl-mediated regulation of LAT-nucleated signaling complexes. Mol Cell Biol 27, 8622–8636 (2007). 10.1128/MCB.00467-07

18 Balagopalan, L. et al. Enhanced T-cell signaling in cells bearing linker for activation of T-cell (LAT) molecules resistant to ubiquitylation. Proc Natl Acad Sci U S A 108, 2885–2890 (2011). 10.1073/pnas.1007098108

19 Rao, N., Dodge, I. & Band, H. The Cbl family of ubiquitin ligases: critical negative regulators of tyrosine kinase signaling in the immune system. J Leukoc Biol 71, 753–763 (2002).

20 Gaud, G., Lesourne, R. & Love, P. E. Regulatory mechanisms in T cell receptor signalling. Nat Rev Immunol 18, 485–497 (2018). 10.1038/s41577-018-0020-8

21 Hofinger, E. & Sticht, H. Multiple modes of interaction between Lck and CD28. J Immunol 174, 3839–3840; author reply 3840 (2005). 10.4049/jimmunol.174.7.3839-a

22 Liaunardy-Jopeace, A., Murton, B. L., Mahesh, M., Chin, J. W. & James, J. R. Encoding optical control in LCK kinase to quantitatively investigate its activity in live cells. Nat Struct Mol Biol 24, 1155–1163 (2017). 10.1038/nsmb.3492

23 Philipsen, L. et al. De novo phosphorylation and conformational opening of the tyrosine kinase Lck act in concert to initiate T cell receptor signaling. Sci Signal 10 (2017). 10.1126/scisignal.aaf4736

24 Casas, J. et al. Ligand-engaged TCR is triggered by Lck not associated with CD8 coreceptor. Nat Commun 5, 5624 (2014). 10.1038/ncomms6624

25 Acuto, O. & Michel, F. CD28-mediated co-stimulation: a quantitative support for TCR signalling. Nat Rev Immunol 3, 939–951 (2003). 10.1038/nri1248

26 Bunnell, S. C. et al. Persistence of cooperatively stabilized signaling clusters drives T-cell activation. Mol Cell Biol 26, 7155–7166 (2006). 10.1128/MCB.00507-06

27 Bunnell, S. C., Barr, V. A., Fuller, C. L. & Samelson, L. E. High-resolution multicolor imaging of dynamic signaling complexes in T cells stimulated by planar substrates. Sci STKE 2003, PL8 (2003). 10.1126/stke.2003.177.pl8

28 Bunnell, S. C. et al. T cell receptor ligation induces the formation of dynamically regulated signaling assemblies. J Cell Biol 158, 1263–1275 (2002). 10.1083/jcb.200203043

29 Campi, G., Varma, R. & Dustin, M. L. Actin and agonist MHC-peptide complex-dependent T cell receptor microclusters as scaffolds for signaling. J Exp Med 202, 1031–1036 (2005). 10.1084/jem.20051182

30 Gioia, L., Siddique, A., Head, S. R., Salomon, D. R. & Su, A. I. A genome-wide survey of mutations in the Jurkat cell line. BMC Genomics 19, 334 (2018). 10.1186/s12864-018-4718-6

31 Cefai, D. et al. CD28 receptor endocytosis is targeted by mutations that disrupt phosphatidylinositol 3-kinase binding and costimulation. J Immunol 160, 2223–2230 (1998).

32 Shan, X. et al. Deficiency of PTEN in Jurkat T cells causes constitutive localization of Itk to the plasma membrane and hyperresponsiveness to CD3 stimulation. Mol Cell Biol 20, 6945–6957 (2000). 10.1128/MCB.20.18.6945-6957.2000

33 Chuang, E. et al. The CD28 and CTLA-4 receptors associate with the serine/threonine phosphatase PP2A. Immunity 13, 313–322 (2000). 10.1016/s1074-7613(00)00031-5

34 Schneider, H., Prasad, K. V., Shoelson, S. E. & Rudd, C. E. CTLA-4 binding to the lipid kinase phosphatidylinositol 3-kinase in T cells. J Exp Med 181, 351–355 (1995). 10.1084/jem.181.1.351

35 Perkins, D. et al. Regulation of CTLA-4 expression during T cell activation. J Immunol 156, 4154–4159 (1996).

36 Hermiston, M. L., Xu, Z. & Weiss, A. CD45: a critical regulator of signaling thresholds in immune cells. Annu Rev Immunol 21, 107–137 (2003). 10.1146/annurev.immunol.21.120601.140946

37 Palacios, E. H. & Weiss, A. Function of the Src-family kinases, Lck and Fyn, in T-cell development and activation. Oncogene 23, 7990–8000 (2004). 10.1038/sj.onc.1208074

38 Balagopalan, L. et al. Plasma membrane LAT activation precedes vesicular recruitment defining two phases of early T-cell activation. Nat Commun 9, 2013 (2018). 10.1038/s41467-018-04419-x

